# Fibroblast State Reversal By MBNL1-Dependent Transcriptome Modification Regulates Cardiac Repair

**DOI:** 10.1101/2021.01.26.428279

**Authors:** Darrian Bugg, Ross Bretherton, Kylie Beach, Anna Reese, Jagadambika Gunaje, Galina Flint, Cole A. DeForest, April Stempien-Otero, Jennifer Davis

## Abstract

Dynamic fibroblast state transitions are responsible for the heart’s fibrotic response to injury, raising the possibility that tactical control of these transitions could alter maladaptive fibrotic outcomes. Transcriptome maturation by the RNA binding protein Muscleblind Like 1 (MBNL1) has emerged as a potential driver of differentiated cell states. Here genetic lineage tracing of myofibroblasts in the injured heart demonstrated that gains in MBNL1 function corresponded to profibrotic fibroblast states. Similarly, in mice cardiac fibroblast specific MBNL1 overexpression induced a transcriptional myofibroblast profile in healthy cardiac fibroblasts that prevented the fibroproliferative phase of cardiac wound healing. By contrast loss of MBNL1 reverted cardiac fibroblasts to a pro-proliferative epicardial progenitor state that limited cardiac fibrosis following myocardial infarction. This progenitor state transition was associated with an MBNL1-dependent destabilization of the mesenchymal transition gene, *Sox9.* These findings suggest that MBNL1 regulation of the fibroblast transcriptome drives state transitions underlying cardiac fibrosis and repair.

## INTRODUCTION

A central obstacle in heart disease is the replacement of healthy muscle with fibrotic scarring. While scarring prevents cardiac rupture following myocardial infarction (MI), it also causes hemodynamic dysfunction and arrhythmias, which rapidly progress the heart towards failure (Gianluigi Savarese and Lars H Lund, 2017). The development of fibrosis results from dynamic state changes in resident cardiac fibroblasts of the *Tcf21* and *Pdfgrα* lineage that first become proliferative and then transition into myofibroblasts, which are defined by the expression of Periostin (Postn) and α-smooth muscle actin (αSMA) along with enhanced production of extracellular matrix (ECM) proteins like Collagens 1 and 3 as well as Fibronectin (Davis and Molkentin, 2014; Dobaczewski et al., 2012; Stempien-Otero et al., 2016). Evidence from murine lineage reporters suggests the myofibroblast state is transient and partially reversible following exposure to pro-fibrotic chemical stimuli by yet unresolved molecular regulatory mechanisms (Kanisicak et al., 2016).

A potential control point for these state transitions is transcriptome maturation, which is mediated by RNA binding proteins like Muscleblind Like 1 (MBNL1). MBNL1 has been shown to modulate a cell’s transcriptional landscape by properly stabilizing, splicing, polyadenylating, and localizing its target mRNAs (Batra et al., 2014; Pascual et al., 2006; Wang et al., 2012). Notably, dysfunctional MBNL1 can cause myotonic dystrophy in addition to differentiation defects during erythrocyte and myofibroblast development (Cheng et al., 2014; Davis et al., 2015). In fibroblasts, we previously identified that myofibroblast state transitions require MBNL1 stabilization of transcripts encoding key signaling factors like serum response factor (Srf) and Calcineurin (CnA), which promote the myofibroblast state, and in the heart myofibroblast density and scarring is reduced in global MBNL1 knockout mice following ischemic injury (Davis et al., 2015). These data prompted the examination of whether targeted modulation of MBNL1 expression *in vivo* could control cardiac fibroblast state transitions and hence fibrotic scarring.

Here Periostin *(Postn)* lineage tracing studies revealed that MBNL1 is upregulated in myofibroblasts but not in quiescent fibroblast progenitors, and similarly MBNL1 is overexpressed in fibroblasts derived from failing human heart biopsies demonstrating the generalizability of this mechanism to human heart disease. Injury-induced cardiac fibroblast state transitions were also analyzed in conjunction with fibrosis following targeted gain or loss of MBNL1 function in mice. In the absence of injury cardiac fibroblast specific MBNL1 overexpression inhibited proliferation and induced a myofibroblast transcriptional profile. Despite the presence of myofibroblasts in MBNL1 transgenic mice, myocardial fibrosis was not detectable at baseline but accentuated with an additive injury stimulus. Rather than forming myofibroblasts following MI, mice with targeted deletion of MBNL1 in resident cardiac fibroblasts reverted to a highly proliferative epicardial progenitor state marked by the overexpression of *Aldh1a2* and downregulation of several epicardial to mesenchymal transition (EMT) genes like *Sox9* and *Tcf21* that underlie fibroblast specification during cardiac development. Importantly this fibroblast state reversion reduced the magnitude of fibrotic scarring and limited maladaptive dilated myocardial remodeling.

## RESULTS

### MBNL1 Is Upregulated In Mouse and Human Cardiac Myofibroblasts

Previous work has shown that cardiac fibroblasts undergo three state changes during the course of infarct repair including: a proliferative state in which resident fibroblasts expand (days 1-4); a myofibroblast state that forms provisional matrix and contracts the wound (days 4-14), and a matrifibrocyte state (days 14-21) in which osteogenic and cartilage genes are expressed to mature the ECM (Daseke et al., 2020; Fu et al., 2018; Mouton et al., 2019a). Here the transcriptional profile of cardiac fibroblasts was examined following myocardial infarction (MI) by RNAseq. Principal component analysis (PCA) of these data demonstrated that at each of these transitional time points cardiac fibroblasts occupy distinct positions in compressed transcriptional space, which is consistent with the notion that cardiac fibroblasts undergo multiple state transitions following MI (**Fig. 1A**). Here most of the variance between sham and injured cardiac fibroblasts was observed in the first component, while shifts in the second component appeared dependent on the amount of time after injury (**Fig. 1A**). The heatmap in **Fig. 1B** depicts 5,141 genes that were differentially expressed in cardiac fibroblasts over the duration of infarct repair. These genes were functionally clustered using Gene Ontology (GO) analysis as denoted by the colored bars and numbers. Genes involved in developmental functions were downregulated throughout the course of MI (**Fig. 1C**); whereas, genes associated with the ECM were upregulated at days 4 and 14 (**Fig. 1D**), tracking with previous data showing the expression of provisional matrix followed by ECM maturation (Fu *et al.,* 2018). In addition, cell cycle genes were highly expressed at day 4 following MI and then down regulated for the remainder of the time course, which matches the timing of the fibroproliferative phase of infarct repair (**Fig. 1E**).

**Figure 1:**
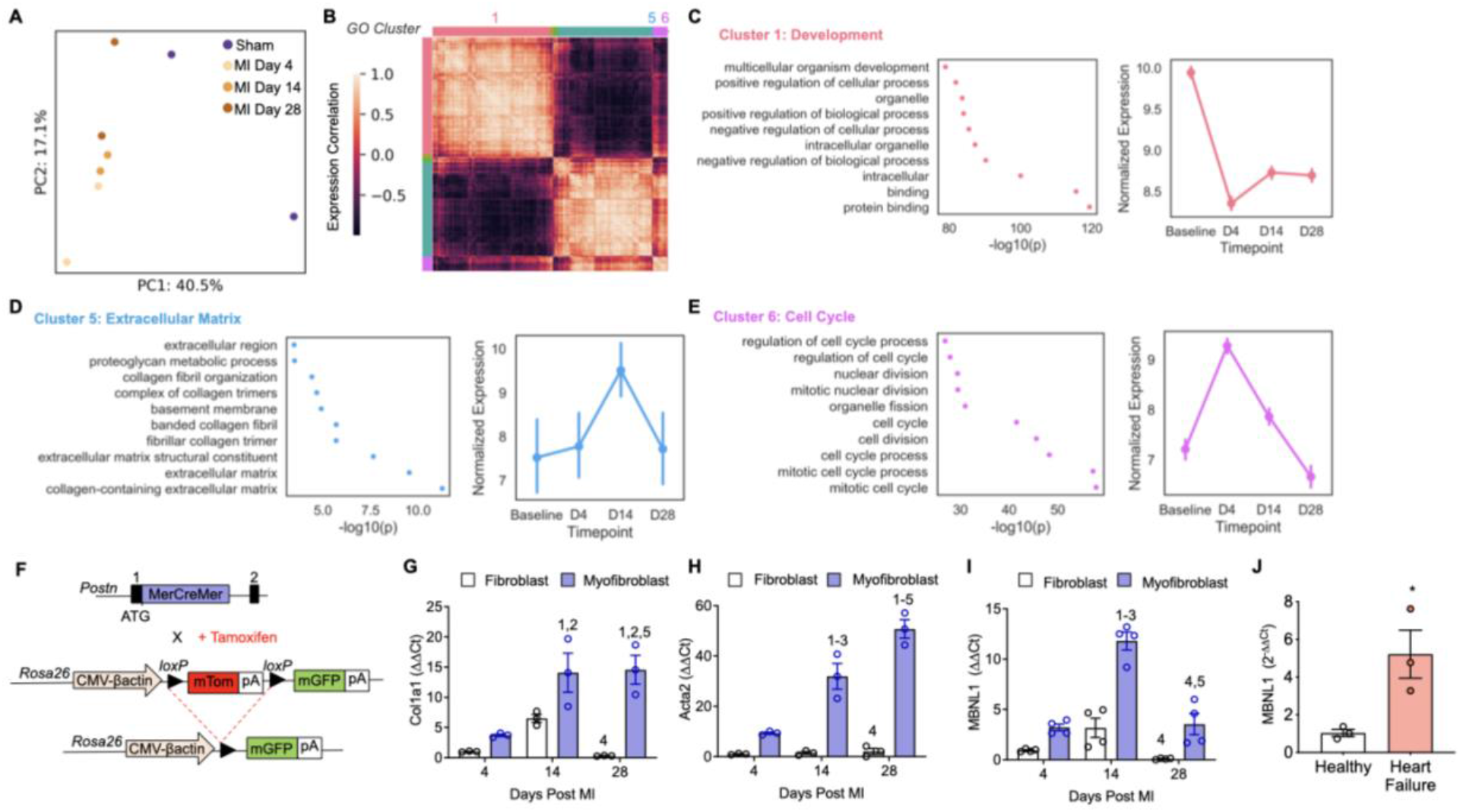
MBNL1 upregulation coincides with fibroblast activation following myocardial infarction. **(A)** PCA analysis of flow-sorted MEFSK4 sorted cardiac fibroblasts from sham and 4, 14, and 28 days post-MI demonstrating a time and injury-dependent change in fibroblast state. **(B)** Pairwise correlation heatmap of 5,141 genes with significant differential expression between timepoints, color coded by gene ontology (GO) functional clusters where 1 (Salmon) denotes Development, 5 (Light Blue) denotes ECM, and 6 (Pink) denotes Cell Cycle. **(C-E)** GO functional enrichment analysis **(left, panel)** and mean±95%CI expression of major gene clusters across infarct repair **(right, panel)**. **(F)** Schematic depicting myofibroblast lineage reporter mouse model generation: **(top)** Tamoxifen-inducible MerCreMer (MCM) *Postn* knock-in is crossed with **(bottom)** mice expressing a membrane-targeted dual fluorescent reporter (mT/mG) driven by a CMV-βactin promoter from the *Rosa26* locus. Fold change in **(G)** *Col1a1,* **(H)** *Acta2,* and **(I)** *Mbnl1* gene expression in flow sorted cardiac myofibroblasts (MEFSK4^+^, mG^+^) and fibroblasts (MEFSK4^+^) relative to day 4 fibroblast (MEFSK4^+^) group. Gene expression was normalized to 18s expression and fold change calculated using the ΔΔCt method, dots represent biological replicates, n=3 mice in each condition and bars represent mean±SEM and ANOVA statistical test with Tukey post hoc comparisons. Numbers represent significant pairwise comparisons at p<0.05: **1**-4d Fibroblast, **2**-4d Myofibroblast, **3**-14d Fibroblast, **4**-14d Myofibroblast, **5**-28d Fibroblast, **6**-28d Myofibroblast. **(J)** Fold change in *MBNL1* gene expression in fibroblasts isolated from failing versus healthy human hearts. Gene expression for each sample was first normalized to *GAPDH* expression and then fold change calculated using the ΔΔCt method. Dots represent biological replicates (n=3 per condition) and bars represent mean±SEM, unpaired t-test with *p<0.05.

Since previous studies have demonstrated heterogeneity within the cardiac fibroblast population following MI (Farbehi et al., 2019; Skelly et al., 2018), *Postn* lineage reporter mice (Postn^iCre^-mT/mG) were used to label and then segregate activated myofibroblasts from the total fibroblast population by flow cytometry to compare gene expression profiles between the two groups over the course of infarct repair. These lineage reporter mice were generated by crossing mice containing a tamoxifen (Tam)-inducible Cre recombinase knocked into the *Postn* locus (Postn^iCre^) with mice containing a ubiquitously expressed dual fluorescent reporter (mT/mG, **Fig. 1F**). With this model Tam-induced cells expressing *Postn*, an early myofibroblast marker (Kanisicak et al., 2016), are labeled with membrane-targeted green fluorescent protein (mG) (**Fig. 1F**). For this experiment Tam-labeling began at the time of surgery and mice remained on Tam for 4, 14, or 28 days following MI. Real-time polymerase chain reaction (RT-PCR) analysis of the *Postn*-traced fibroblasts showed a significant upregulation of core myofibroblast genes like *Col7a7* and *Acta2* relative to quiescent fibroblasts at all time points (**Fig1. G-H**). Owing to our previous work identifying MBNL1 as an activator of myofibroblast differentiation (Davis et al., 2015), MBNL1 gene expression was also examined. In *Postn*-traced myofibroblasts MBNL1 expression increased 3-fold relative to quiescent fibroblasts 4 days following MI (**Fig. 1I**). By day 14 MBNL1 expression more than tripled but regressed back to quiescent levels by day 28 (**Fig. 1I**). To examine whether this upregulation in MBNL1 is generalizable to human heart failure, cardiac fibroblasts were isolated from left ventricular biopsies taken from patients receiving left ventricular assist devices-a human cardiac tissue source rich in fibroblasts that have transitioned to a myofibroblast state (Farris et al., 2017). Relative to healthy human fibroblasts there was a significant increase in MBNL1 expression in failing cardiac fibroblasts (**Fig. 1J**), suggesting that dynamic changes in fibroblast specific MBNL1 expression are associated with the timing and expression of transcripts responsible for a myofibroblast phenotype in both mice and humans.

### MBNL1 Expression Induces A Myofibroblast Transcriptional Profile In Resident Cardiac Fibroblasts

The upregulation of MBNL1 in coordination with core myofibroblast genes during myocardial infarct repair suggests that MBNL1 plays a role in myofibroblast state determination. To examine whether MBNL1 overexpression is sufficient for transitioning cardiac fibroblasts into myofibroblasts, conditional MBNL1 transgenic (Tg) mice were crossed with mice containing a cardiac fibroblast specific Cre driver, which was engineered by knocking a Tam-inducible Cre recombinase into the *Tcf21* locus (Tcf21^iCre^, **Fig. 2A**, (Acharya et al., 2011,2012)). At 6 weeks of age non-transgenic (NTG-Tcf21^iCre^) and MBNL1 Tg-Tcf21^iCre^ littermates were administered Tam for 2-weeks at which time cardiac fibroblasts were isolated, purified by flow cytometry, and subjected to RNAseq analysis that identified 420 differentially expressed genes between groups (**Fig. 2B**). Of those differentially expressed genes many were previously identified as MBNL1 regulatory targets in an RNA immunoprecipitation RNAseq analysis (RIPseq) (**Supplemental Fig. 1,** (Davis et al., 2015)). Here MBNL1-bound genes like *Lox, Postn, Fn1,* and *Ccn4* were all upregulated in MBNL1 Tg-Tcf21^iCre^ fibroblasts (**Supplemental Fig. 1**). Notably, all of these genes have been associated with myofibroblast differentiation and fibrosis (Kanisicak et al., 2016; Martin et al., 2020; Ono et al., 2018; Sun et al., 2020). Principal component analysis of the RNAseq data set revealed that at baseline fibroblast specific expression of MBNL1 causes the phenotype to transition into the same transcriptional space as non-transgenic fibroblasts subjected to MI (**Fig. 2C**). Given that days 4-14 represents the time frame for maximal myofibroblast differentiation, these data provide further evidence that MBNL1 expression is sufficient to transition the cardiac fibroblast transcriptome toward a myofibroblast phenotype in an uninjured heart. Gene ontology (GO) term analysis of differentially expressed genes revealed there was a significant enrichment in gene categories for ECM and several cardiovascular developmental pathways, which fits with a transgene-dependent induction of a myofibroblast transcriptional profile (**Fig. 2D**). Further examination of this RNAseq data set demonstrated that several fundamental transcriptional markers of the myofibroblast cell state like *Acta2, Postn, Fn1, Lox,* and *Col5* were upregulated in uninjured MBNL1 Tg-Tcf21^iCre^ fibroblasts when compared to non-transgenic littermates (**Fig. 2E**). While just 2 weeks of MBNL1 overexpression transitions the fibroblast transcriptome to the transcriptional state space of a profibrotic myofibroblast (**Fig. 2C & E**), no differences were identified in organ level fibrosis (**Fig. 2F-G**), myocardial αSMA^+^ myofibroblasts (**Fig. 2H-I**), or cardiac structure (**Fig. 2J**) and function (**Fig. 2K**) when comparing MBNL1 Tg-Tcf21^iCre^ and NTG-Tcf21^iCre^ groups.

It is possible that an additional pathologic stimulus like injury or enhanced myocardial stiffness is needed to achieve the myofibroblast phenotype at a proteomic and physiologic level, so here cardiac fibroblasts were isolated from non-transgenic and MBNL1 Tg-Tcf21^iCre^ mice and immunofluorescent imaging was used to quantify the number of cardiac fibroblasts that form αSMA^+^ stress fibers, a morphologic marker of myofibroblasts (**Fig. 2L**) (Tallquist and Molkentin, 2017). Without any pharmacologic induction there was a 2-fold increase in the number of fibroblasts with αSMA^+^ stress fibers in the MBNL1 Tg-Tcf21^iCre^ group relative to NTG-Tcf21^iCre^. Moreover, inducing myofibroblast differentiation with a known profibrotic agonist, transforming growth factor-β (TGFβ), failed to additively increase the number of MBNL1 Tg-Tcf21^iCre^ fibroblasts with αSMA^+^ stress fibers, which differs from the significant transformation of NTG-Tcf21^iCre^ fibroblasts into myofibroblasts with TGFβ treatment. These data indicate that MBNL1 can induce myofibroblast state at the proteomic and physiologic levels with the appropriate cues as well as regulate components of the TGFβ pathway. Previously we demonstrated that one of those components is serum response factor (SRF), which is stabilized by MBNL1 and required for TGFβ-dependent myofibroblast differentiation (Davis et al., 2015). Here acute shRNA knockdown of SRF blocked TGFβ-dependent myofibroblast differentiation in both MBNL1 Tg-Tcf21^iCre^ and NTG-Tcf21^iCre^ fibroblasts demonstrating that MBNL1 requires SRF function to promote the myofibroblast phenotype.

Given that MBNL1 Tg-Tcf21^iCre^ fibroblasts failed to show statistically significantly gains in the number of αSMA^+^ myofibroblasts or ECM quantity in the heart at baseline but had the phenotype emerge at the physiologic level *in vitro* when activated by stiff tissue culture plastic, MIs were surgically applied to MBNL1 Tg-Tcf21^iCre^ and NTG-Tcf21^iCre^ mice two weeks following MBNL1 transgene induction. Despite the expression of myofibroblasts genes in the fibroblasts of MBNL1 Tg-Tcf21^iCre^ mice (**Fig. 2E**), only a modest increase in myofibroblast density and overall fibrosis was observed in response to MI (**Fig. 3A-D**). A possible explanation for the unexpected modest change in myofibroblast number and quantity of fibrosis is that the baseline myofibroblast transcriptional profile in MBNL1 Tg-Tcf21^iCre^ cardiac fibroblasts negatively impacted the fibroproliferative phase of MI-induced wound healing, which occurs prior to their transition to a myofibroblast state (Fu et al., 2018). Because defective proliferation could limit the number of myofibroblast progenitors and hence the fibrotic response to injury, we sought to examine this hypothesis. Here MBNL1 Tg-Tcf21^iCre^ and NTG-Tcf21^iCre^ littermates were Tam-induced, subjected to MI, and given 5-ethynyl-2’-deoxyuridine (EdU) to assess cell proliferation in myocardial sections 2-days after injury. Here the colocalization of EdU^+^ (Click-iT) and PDGFRα^+^ was used to identify proliferating fibroblasts in myocardial sections (**Fig. 3E-F**). MBNL1 Tg-Tcf21^iCre^ mice had approximately 20% fewer EdU^+^ fibroblasts in comparison to NTG-Tcf21^iCre^ controls (**Fig. 3F**). To confirm that MBNL1 causes a proliferation defect, cardiac fibroblasts were isolated from MBNL1 Tg-Tcf21^iCre^ and NTG-Tcf21^iCre^ mice and put into primary culture where EdU incorporation was examined under differentiation conditions (**Fig. 3G,** 2% Serum). Similar to the *in vivo* findings MBNL1 Tg-Tcf21^iCre^ fibroblasts had significantly reduced proliferation capacity (**Fig. 3G**). Notably, this proliferation defect was also confirmed by RNAseq analysis of cardiac fibroblasts isolated from MBNL1 Tg-Tcf21^iCre^ and NTG-Tcf21^iCre^ mice 4 days after MI in which key positive cell cycle regulators like *Ccnb2, Ccna2, Birc5,* and *Kif23* were all down regulated while cell cycle inhibitors like *Cxcl1* and *Cdkn1a* were significantly upregulated (**Fig. 3H**). Collectively, these data suggest that despite the transcriptional myofibroblast phenotype, decreased fibroblast proliferation in this model inhibited any additive effects on myofibroblast density and cardiac scarring. Unexpectedly, these molecular state changes in cardiac fibroblasts did elicit a 10% drop in fractional shortening and increased left ventricular dilation (LV diameter, **Fig. 3I-J**) suggesting that these state changes still impact maladaptive cardiac remodeling.

**Figure 2:**
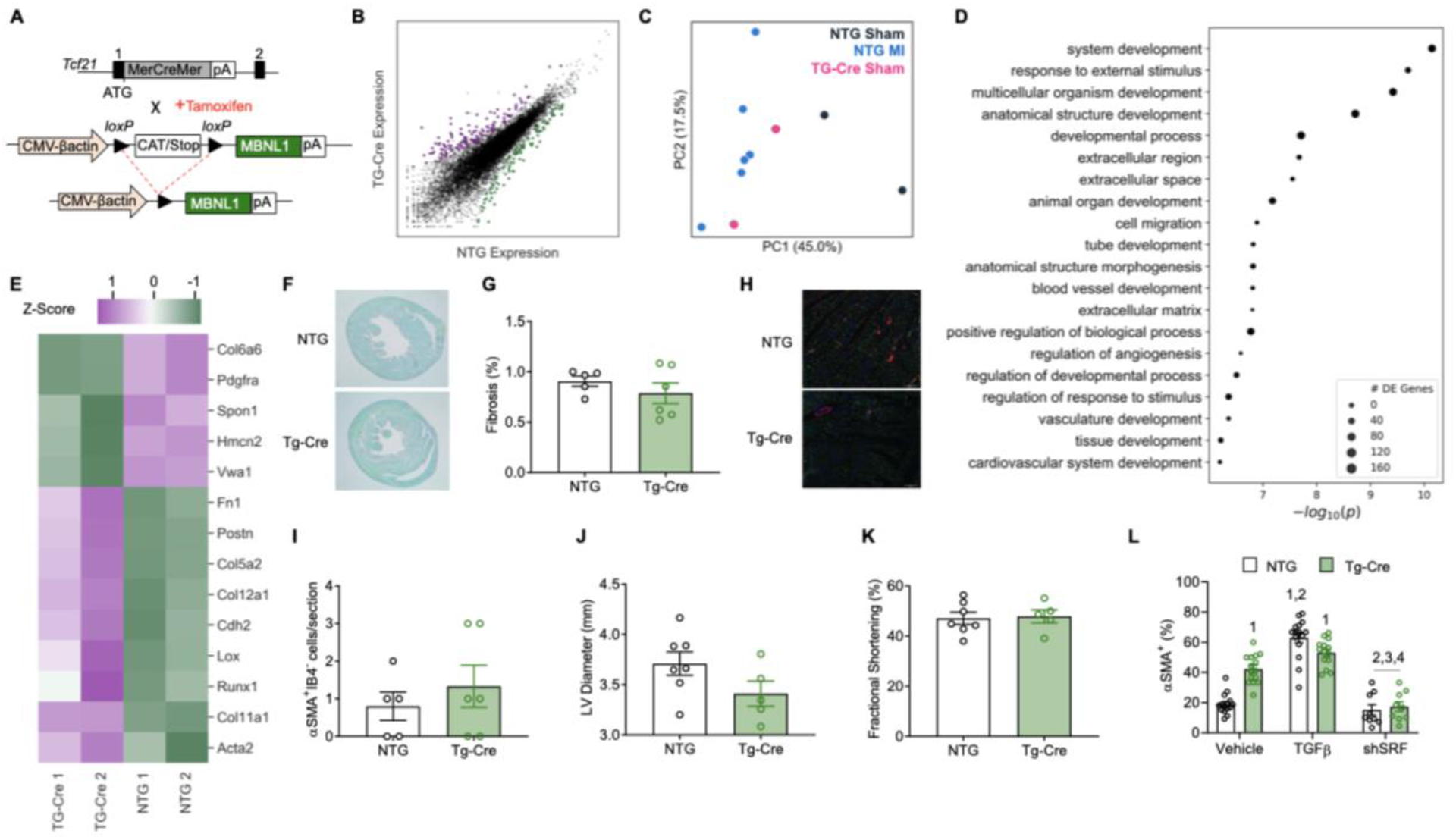
Cardiac fibroblast specific MBNL1 overexpression induces a myofibroblast phenotype at baseline. **(A)** Schematic depicting MBNL1 Tg-Tcf21^iCre^ mouse model generation: **(top)** Tamoxifen-inducible MerCreMer (MCM) *Tcf21* knock-in were crossed with **(bottom)** mice containing a LoxP-flanked Stop cassette followed by a MBNL1 transgene expressed by a CMV-βactin promoter. **(B)** Scatterplot of differentially expressed genes between MEFSK4 sorted cardiac fibroblasts from MBNL1 Tg-Tcf21^iCre^ versus NTG-Tcf21^iCre^. Purple dots represent genes significantly upregulated in MBNL1 Tg-Tcf21^iCre^, green dots represent genes significantly down regulated in MBNL1 Tg-Tcf21^iCre^, black dots represent genes lacking significant differences in expression. **(C)** PCA of MEFSK4 sorted cardiac fibroblasts from sham MBNL1 Tg-Tcf21^iCre^ and NTG-Tcf21^iCre^ mice as well as injured NTG-Tcf21^iCre^ mice. **(D)** Functional gene clustering of differentially expressed genes between MBNL1 Tg-Tcf21^iCre^ and NTG-Tcf21^iCre^ MEFSK4 sorted cardiac fibroblasts at baseline. Number of differentially expressed (DE) genes is correlated with the size of circle in each cluster. **(E)** Heatmap of differentially expressed genes in MBNL1 Tg-Tcf21^iCre^ and NTG-Tcf21^iCre^ MEFSK4 sorted cardiac fibroblasts at baseline. **(F)** Histological images and **(G)** quantification of Sirius Red/Fast Green-stained fibrosis (red) and muscle (green) in MBNL1 Tg-Tcf21^iCre^ (n=6), NTG-Tcf21^iCre^ (n=5) hearts after 2 weeks of Tam administration. Dots represent biological replicates and bars represent mean±SEM and unpaired t-tests with p=0.3551. **(H)** Immunofluorescent imaging and **(I)** quantification of myofibroblasts in the left ventricle of myocardial sections from mice after 14 days of tamoxifen administration. Myofibroblasts are αSMA (red) positive and negative for the endothelial marker isolectinB4 (IB4, green). Nuclei are stained blue. Scale bar=50μM. Dots represent biological replicates MBNL1 Tg-Tcf21^iCre^ (n=6), NTG-Tcf21^iCre^ (n=5) and bars represent mean±SEM and unpaired t-tests with p=0.4675. Echocardiographic analysis of MBNL1 Tg-Tcf21^iCre^ (n=5) and NTG-Tcf21^iCre^ (n=7) mice quantifying **(J)** left ventricle inner diameter at diastole p=0.0828 and **(K)** fractional shortening p=0.8372. Dots represent biological replicates and bars represent mean±SEM, unpaired t-tests used for statistical comparison. **(L)** Quantification of the number of cardiac fibroblasts with αSMA^+^ stress fibers as a percentage of the total number counted. Here cardiac fibroblasts were obtained from MBNL1 Tg-Tcf21^iCre^ and NTG-Tcf21^iCre^ mice and some groups were treated with TGFβ or adenovirally transduced with SRF shRNA. Dots represent biological replicates (n=18), and bars represent mean±SEM. Numbers represent p<0.05 from ANOVA statistical test with Tukey post hoc comparisons. **1**-NTG Vehicle, **2**-Tg-Cre Vehicle, **3**-NTG TGFβ, **4**-Tg-Cre TGFβ, **5**-NTG shSRF, **6**-Tg-Cre shSRF.

**Figure 3:**
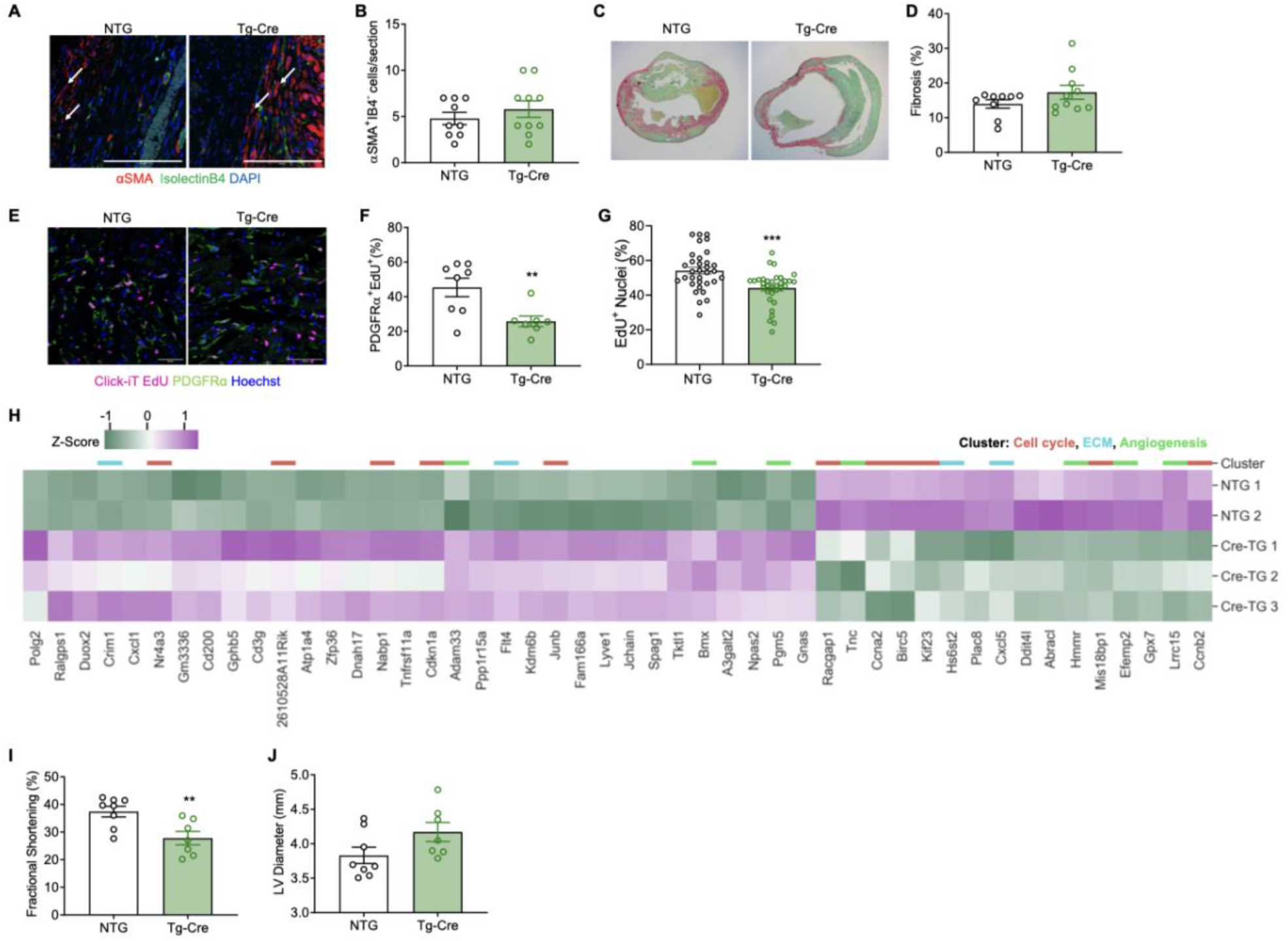
Overexpression of MBNL1 in cardiac fibroblasts prevents MI-dependent fibroblast proliferation. **(A)** Immunofluorescent imaging and **(B)** quantification of myofibroblasts in the border zone of myocardial sections from mice 14 days following MI. Myofibroblasts are αSMA (red) positive and negative for the endothelial marker isolectinB4 (IB4, green). Nuclei are stained blue. Arrows show αSMA^+^IB4^-^ cells. Scale bar=50μM. Dots represent biological replicates MBNL1 Tg-Tcf21^iCre^ (n=10), NTG-Tcf21^iCre^ (n=9) and bars represent mean±SEM and unpaired t-tests, p=0.3791. **(C)** Histological images and **(D)** quantification of Sirius Red/Fast Green-stained fibrosis (red) and muscle (green) in MBNL1 Tg-Tcf21^iCre^ (n=10), NTG-Tcf21^iCre^ (n=9) hearts. Unpaired t-tests with p=0.1808. **(E)** Immunofluorescent staining and **(F)** quantification of proliferating fibroblasts 2 days post MI in MBNL1 Tg-Tcf21^iCre^ (n=8) and NTG-Tcf21^iCre^ (n=7) hearts. Proliferating fibroblasts are co-stained with PDGFRα (green) and EdU (pink). Nuclei are stained blue. Arrows show PDGFRα^+^,EdU^+^ fibroblasts. Scalebar=50μM. Dots represent biological replicates and bars represent mean±SEM and unpaired t-tests,**p<0.01. **(G)** Quantification of proliferating fibroblasts *in vitro* with low serum conditions. MBNL1 Tg Tcf21^iCre^ (n=11 biological replicates), NTG-Tcf21^iCre^ (n=11 biological replicates). Dots represent both biologic and technical replicates. Bars represent mean+SEM, unpaired t-test, ***p<0.001. **(H)** Heatmap of differentially expressed genes in MBNL Tg-Tcf21^iCre^ versus NTG-Tcf21^iCre^ MEFSK4 sorted fibroblasts 14 days post MI. Colored bars represent functionally clustered genes: Extracellular matrix (ECM) = blue, Cell cycle = red, and Angiogenesis = green. Echocardiographic analysis of MI-injured MBNL1 Tg-Tcf21^iCre^ (n=7) and NTG-Tcf21^iCre^ (n=8) mice for **(I)** ventricular fractional shortening **p<0.01 and **(J)** left ventricle inner diameter during diastole p=0.0828. Dots represent biological replicates and bars represent mean±SEM, unpaired t-tests for statistical comparison.

### Cardiac Fibroblast Specific Deletion of MBNL1 Blocks Myofibroblast Differentiation & Cardiac Fibrosis

To examine the requirement for MBNL1 in cardiac fibroblast to myofibroblast state transitions, conditional MBNL1 knockout mice (MBNL1^Fl/Fl^) were generated and then crossed with Tcf21^iCre^ mice (**Fig. 4A**). Western blot of TGFβ-treated cardiac fibroblasts obtained from Tam-induced MBNL1^Fl/Fl^ and MBNL1^Fl/Fl^-Tcf21^iCre^ mice demonstrated that nearly complete deletion of MBNL1 is achieved in cardiac fibroblast specific MBNL1 knockouts (**Fig. 4B**). In the absence of injury loss of MBNL1 caused no observable physiological changes by echocardiography (*data not shown)* but following injury MBNL1^Fl/Fl^-Tcf21^iCre^ mice had reduced ventricular dilation in comparison to MBNL1^Fl/Fl^ littermates (**Fig. 4C**). Neither group exhibited differences in systolic function (**Fig. 4D**). In addition, MBNL1^Fl/Fl^-Tcf21^iCre^ mice had a 40% reduction in both fibrosis and myofibroblast density when compared to MBNL1^Fl/Fl^ controls suggesting that MBNL1 is required for myofibroblast formation and cardiac fibrosis (**Fig. 4E-H**). To confirm the reduction in fibrotic scarring and myofibroblast density was due to myofibroblast differentiation defects in the MBNL1^Fl/Fl^-Tcf21^iCre^ fibroblasts, cardiac fibroblasts were isolated from MBNL1^Fl/Fl^ and MBNL1^Fl/Fl^-Tcf21^iCre^ mice and subjected to RNAseq analysis. In accordance with the *in vivo* histological analysis, MBNL1^Fl/Fl^-Tcf21^iCre^ fibroblasts had downregulated key myofibroblast genes like *Acta2, Col1a1, Col1a2,* and *Postn* (**Fig. 4I**). There was also a reduction in transcripts such as, *Comp* and *Ccn4/5,* which are genes associated with a cell state required for maturing fibrotic ECM (Fu et al., 2018). Notably, *Acta2, Col1a1, Lox, Runx, Postn* and *Ccn4* were all differentially upregulated in MBNL1 Tg-Tcf21^iCre^ fibroblasts at baseline (**Fig. 2E**); whereas, the fibroblast progenitor marker *Pdgfrα* was upregulated in MBNL1 knockout fibroblasts (**Fig. 4I**) but conversely downregulated by MBNL1 overexpression (**Fig. 2E**).

**Figure 4:**
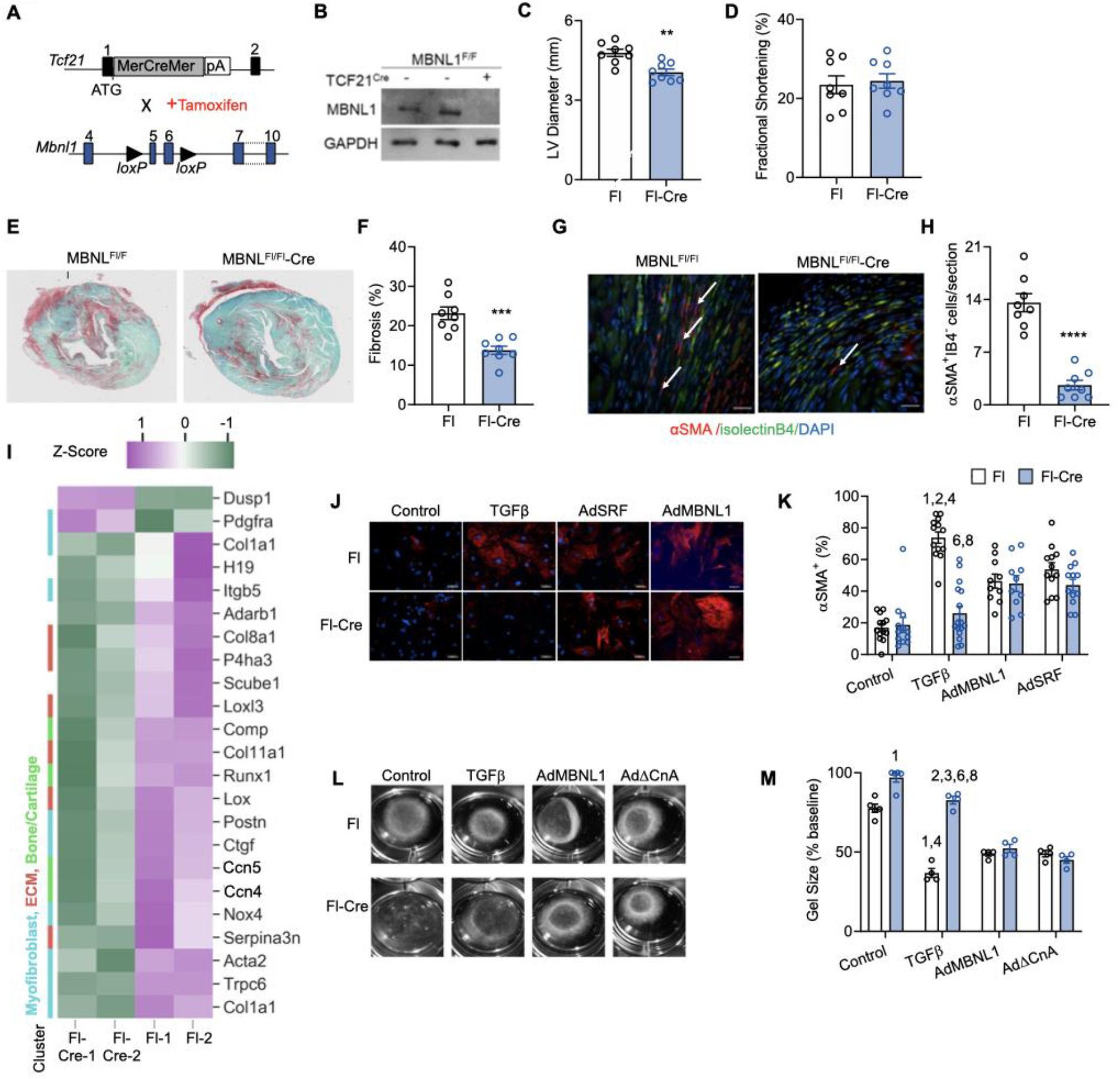
MBNL1 is required for cardiac fibroblast to myofibroblast state transitions and MI-induced fibrotic scarring. **(A)** Schematic depicting the generation of cardiac fibroblast specific MBNL1 knockout mice (MBNL1^Fl/Fl^-Tcf21^iCre^): (**top**) Tam-inducible MerCreMer (MCM) *Tcf21* knock-ins were crossed with (**bottom**) mice homozygous for LoxP-targeted *Mbnl1* alleles (MBNL1^Fl/Fl^). **(B)** Western blot analysis for MBNL1 protein expression from Tam-treated MBNL1^Fl/Fl^-Tcf21^iCre^ and MBNL1^Fl/Fl^ cardiac fibroblasts + recombinant TGFβ. GAPDH was used as a loading control. Echocardiographic analysis of **(C)** left ventricle inner diameter **p<0.01 and **(D)** ventricular fractional shortening p=0.7499 14 days post MI in MBNL1^Fl/Fl^-Tcf21^iCre^ (n=8) and MBNL1^Fl/Fl^ (n=8) mice. Dots represent biological replicates and bars represent mean±SEM, unpaired t-tests for statistical comparison. **(E)** Histological images and **(F)** quantification of Sirius Red/Fast Green-stained fibrosis (red) and muscle (green). Dots represent biological replicates and bars represent mean±SEM, unpaired t-tests,***p<0.001. **(G)** Immunofluorescent imaging and **(H)** quantification of myofibroblasts in the border zone of myocardial sections from mice 14 days following MI. Myofibroblasts are αSMA (red) positive and negative for the endothelial marker isolectinB4 (IB4, green). Nuclei are stained blue. Arrows show αSMA^+^IB4^-^ cells. Scalebar=50μM. MBNL1^Fl/Fl^-Tcf21^iCre^ (n=8), MBNL1^Fl/Fl^ (n=8). Dots represent biological replicates and bars represent mean±SEM, unpaired t-tests, ****p<0.0001. **(I)** Heatmap of differentially expressed genes in MBNL1^Fl/Fl^-Tcf21^iCre^ versus MBNL1^Fl/Fl^ MEFSK4 sorted fibroblasts 14 days post MI. Colored bars represent functionally clustered genes: myofibroblast=blue, ECM=red, and bone/cartilage=green. **(J)** Immunofluorescent images and **(K)** quantification of the number of cardiac fibroblasts with αSMA^+^ stress fibers as a percentage of the total number counted. αSMA^+^ stress fibers (red) and nuclei (blue). Dots represent biological replicates (n=12), and bars represent mean±SEM. Numbers represent p<0.05 from ANOVA statistical test with Tukey post hoc comparisons. **1**-Fl Control, **2**-Fl-Cre Control, **3**-Fl TGFβ, **4**-Fl-Cre TGFβ, **5**-Fl AdMBNL, **6**-Fl-Cre AdMBNL1, **7**-Fl AdSRF, **8**-Fl-Cre AdSRF. **(L)** Representative images and **(M)** quantification of contracted collagen gel matrices seeded with Tam-treated MBNL1^Fl/Fl^-Tcf21^iCre^ and MBNL1^Fl/Fl^ cardiac fibroblasts. Dots represent biological replicates (n=4), and bars represent mean±SEM. Numbers represent p<0.05 from ANOVA statistical test with Tukey post hoc comparisons. **1**-Fl Control, **2**-Fl-Cre Control, **3**-Fl TGFβ, **4**-Fl-Cre TGFβ, **5**-Fl AdMBNL1, **6**-Fl-Cre AdMBNL1, **7**-Fl AdCna, **8**-Fl-Cre AdCnA.

*In vitro* myofibroblast differentiation assays were also used to validate these differentiation defects in MBNL1^Fl/Fl^-Tcf21^iCre^ cardiac fibroblasts. On average 73%±3.46 of MBNL1^Fl/Fl^ control fibroblasts developed αSMA^+^ stress fibers when treated with TGFβ versus MBNL1^Fl/Fl^-Tcf21^iCre^ cardiac fibroblasts that were refractory to the profibrotic treatment showing no change in the number of fibroblasts with developed αSMA stress fibers (**Fig. 4J-K**). This result suggests that MBNL1 is required for TGFβ-mediated fibroblast to myofibroblast state changes. Given MBNL1 stabilizes SRF, it was reasoned that expressing a stabilized version of SRF should rescue the myofibroblast differentiation defect observed in MBNL1^Fl/Fl^-Tcf21^iCre^ cardiac fibroblasts (Davis et al., 2015). To address this hypothesis MBNL1^Fl/Fl^ and MBNL1^Fl/Fl^-Tcf21^iCre^ fibroblasts were adenovirally transduced with SRF (AdSRF), and as predicted TGFβ-mediated myofibroblast differentiation was rescued in MBNL1 knockout fibroblasts (**Fig. 4J-K**). Similarly, adenoviral reexpression of MBNL1 (AdMBNL1) in MBNL1^Fl/Fl^-Tcf21^iCre^ cardiac fibroblasts rescued myofibroblast differentiation (**Fig. 4J-K**), indicating these fibroblasts have the ability to undergo programmed myofibroblast state transitions but require MBNL1 function and its downstream regulatory targets to facilitate the state change.

Another hallmark of the myofibroblast phenotype is a gain in contractile function needed to close wounds. Contractile function was measured using collagen gel contraction assays. Here MBNL1^Fl/Fl^ or MBNL1^Fl/Fl^-Tcf21^iCre^ cardiac fibroblasts were seeded in a collagen gel and 48 hours later gel area was measured as a surrogate for cell contractility. MBNL1^Fl/Fl^ control cardiac fibroblasts contracted gels down to ~80% of their original size; whereas, MBNL1^Fl/Fl^-Tcf21^iCre^ cardiac fibroblasts decreased collagen gel size by only 4% (**Fig. 4L-M**) demonstrating these MBNL1 knockout cardiac fibroblasts have limited to no contractile function. As expected, the addition of TGFβ caused significant contraction of the gels containing MBNL1^Fl/Fl^ cardiac fibroblasts, but those containing MBNL1^Fl/Fl^-Tcf21^iCre^ cardiac fibroblasts were insensitive to TGFβ (**Fig. 4L-M**). Similar to results from the myofibroblast differentiation assays (**Fig. 4J-K**), adenoviral re-expression of MBNL1 (AdMBNL1) or expression of a constitutively active splice variant of calcineurin (AdΔCnA), which is alternatively spliced by MBNL1 and necessary for myofibroblast formation (Davis et al., 2012, 2015), restored the contractile phenotype in MBNL1^Fl/Fl^-Tcf21^iCre^ cardiac fibroblasts demonstrating again that acquiring myofibroblast contractile function requires MBNL1 and its regulatory transcripts (**Fig. 4L-M**).

### Injury Reverts MBNL1-Depleted Cardiac Fibroblasts To A Proliferative Epicardial Progenitor State

While the evidence indicated that MBNL1 knockout cardiac fibroblasts are unable to differentiate into myofibroblasts in response to injury, it was unclear where these cells reside in molecular state space. This question was examined by isolating and purifying cardiac fibroblasts via flow cytometry 4 days after MI and performing RNAseq analysis. Relative to MBNL1^Fl/Fl^ controls MBNL1^Fl/Fl^-Tcf21^iCre^ fibroblasts had a 50% reduction in cardiac fibroblast specification transcripts involved in epicardial to mesenchymal transition (EMT) that include: *Tcf21, Ets2, Tbx20, TGFβ3, Adamts1* and *17* (**Fig. 5A**). By contrast, there was a 2-fold increase in *Aldh1a2* expression (**Fig. 5A**), which is a gene required for epicardial cell development (Xavier-Neto et al., 2000). Together this gene expression analysis suggests the MI triggers MBNL1^Fl/Fl^-Tcf21^iCre^ fibroblasts to dedifferentiate to an epicardial progenitor state.

**Figure 5:**
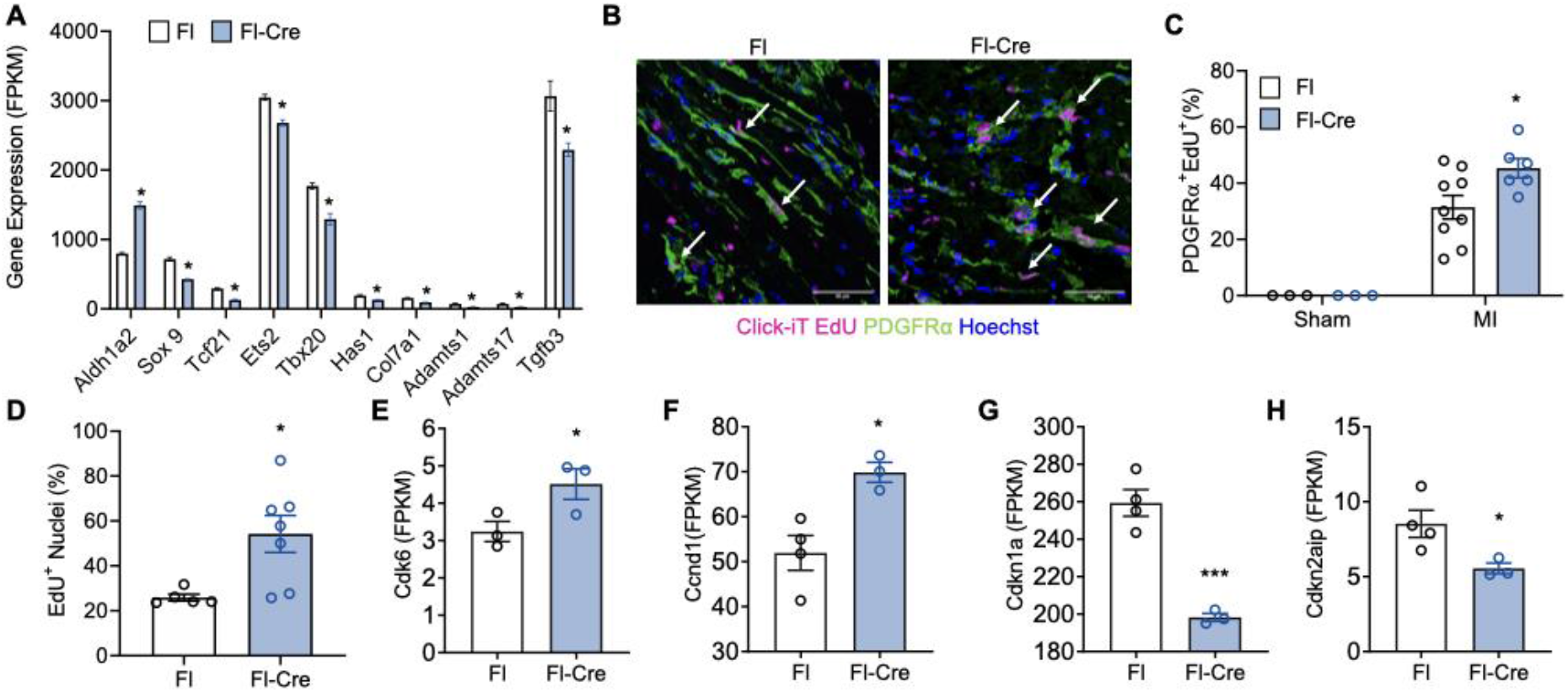
MBNL1 deficiency reverts cardiac fibroblast to an epicardial progenitor state following injury. **(A)** RNAseq analysis of statistically significant (p<0.05) EMT and cardiac fibroblast specification genes expressed in MBNL1^Fl/Fl^ Tcf21^iCre^ (n=4) and MBNL1^Fl/Fl^ (n=6) mice 4 days post MI. Bars represent mean±SEM. **(B-C)** Immunofluorescent staining and quantification of proliferating fibroblasts in myocardial sections from MBNL1^Fl/Fl^-Tcf21^iCre^ (Sham n=3, MI n=6) and MBNL1^Fl/Fl^ (Sham n=3, MI n=9) mice 2 days post MI. Proliferating fibroblasts are PDGFRα^+^ (green) and EdU^+^ (pink). Nuclei are stained blue. Arrows show PDGFRα^+^,EdU^+^ cells. Scalebar=50μM. Dots represent biological replicates and bars represent mean±SEM and unpaired t-tests,*p<0.05. **(D)** Quantification of MBNL1^Fl/Fl^-Tcf21^iCre^ and MBNL1^Fl/Fl^ cardiac fibroblast proliferation *in vitro* with low serum conditions. Dots represent biological replicates (n=7 per genotype) and bars represent mean±SEM, unpaired t-test, *p<0.05. Gene expression (FPKM, fragments per kilobase of transcript) of positive **(E-F)** and **(G-H)** negative cell cycle in MBNL1^Fl/Fl^-Tcf21^iCre^ (n=3) and MBNL1^Fl/Fl^ (n=4) MEFSK4 sorted cardiac fibroblasts 4 days post MI. Dots represent biological replicates and bars represent mean+SEM, unpaired t-test, *p<0.05, ***p<0.001.

Often dedifferentiation is associated with heightened cellular proliferation (Daseke et al., 2020; Fu et al., 2018; Mouton et al., 2019b), and given MBNL1 overexpression inhibited cardiac fibroblast proliferation in MBNL1 Tg-Tcf21^iCre^ mice (**Fig. 3E-G**), we reasoned that loss of MBNL1 function would promote cardiac fibroblast proliferation. MBNL1^Fl/Fl^ and MBNL1^Fl/Fl^-Tcf21^iCre^ were subjected to MI and dosed with EdU to label cells over the first 2 days of MI as described above. In MBNL1^Fl/Fl^-Tcf21^iCre^ hearts 45% of the total PDGFRα^+^ fibroblast population co-labeled with EdU. In comparison only 31% of cardiac fibroblasts were proliferating in MBNL1^Fl/Fl^ controls (**Fig. 5B-C**). This was further validated *in vitro* where a 2-fold increase in EdU^+^ cardiac fibroblasts was observed in MBNL1^Fl/Fl^-Tcf21^iCre^ mice when compared to MBNL1^Fl/Fl^ controls (**Fig. 5D**). The heightened proliferation capacity of MBNL1^Fl/Fl^-Tcf21^iCre^ cardiac fibroblasts was further validated by RNAseq analysis of fibroblasts isolated 4 days after MI. Relative to MBNL1^Fl/Fl^, cardiac fibroblasts from MBNL1^Fl/Fl^-Tcf21^iCre^ mice had increased expression of positive cell cycle regulators *Cdk6* and *Ccnd1* along with decreased expression of potent cell cycle inhibitors *Cdkn7a* and *Cdkn2aip* (**Fig. 5E-H**), both of which were previously shown to be bound and regulated by MBNL1(Davis et al., 2015). Again, there was antithetical expression of *Cdkn1a* and several cyclins in MBNL1 Tg-Tcf21^iCre^ mice (**Fig. 3H**) showing the regulatory power of MBNL1 function on core mediators of both cell proliferation and differentiation. Furthermore, the combination of an MI-dependent downregulation of fibroblast specification genes and enhanced proliferation capacity in MBNL1^Fl/Fl^-Tcf21^iCre^ cardiac fibroblasts suggests these cells have indeed transited back to an early developmental state.

### MBNL1-dependent Stabilization of SOX9 Rescues Myofibroblast Differentiation In MBNL1 Knockout Cardiac Fibroblasts

Still left open is which MBNL1-regulated transcripts are driving these MI-dependent changes in cardiac fibroblast state. To address this question we mined two of our previously published screens that identified MBNL1 regulated transcripts and inducers of myofibroblast differentiation for candidate genes that were (1) bound by MBNL1, (2) induce myofibroblast differentiation, and (3) knocked down in MBNL1^Fl/Fl^ Tcf21^iCre^ cardiac fibroblasts 4 days after MI (Davis et al., 2015). As demonstrated by the Venn diagram only one gene, *Sox9*, met all of the criteria (**Fig. 6A**). *Sox9* is a transcription factor essential for EMT and chondrocyte differentiation that has been previously associated with cardiac myofibroblast differentiation and fibrosis (Lacraz et al., 2017; Scharf et al., 2019). In flow cytometry sorted Postn^+^ myofibroblasts *Sox9* expression increased 8.5-fold relative to its expression in quiescent fibroblasts by 14 days after MI, and it remained elevated for at least a month (**Fig. 6B**). To examine whether *Sox9* is sufficient to promote cardiac fibroblast to myofibroblast differentiation in the absence of MBNL1, cardiac fibroblasts were isolated from MBNL1^Fl/Fl^ and MBNL1^Fl/Fl^-Tcf21^icre^ mice, transfected with *Sox9*, and the number of fibroblasts that transitioned to an myofibroblast phenotype was determined by measuring the percentage of the population with αSMA^+^ stress fibers. Both MBNL1^Fl/Fl^ and MBNL1^Fl/Fl^-Tcf21^iCre^ cardiac fibroblasts had a 2-fold increase in the number of cells with αSMA stress fibers with *Sox9* overexpression (**Fig. 6C**). While it is known that *Sox9* is bound by MBNL1 in fibroblasts (Davis et al., 2015; Girardot et al., 2018), how MBNL1 matures this transcript is still unknown. Since *Sox9* is upregulated in coordination with MBNL1 expression in MI-injured cardiac fibroblasts (**Fig. 1I & 6B**), it was hypothesized that MBNL1 is required for stabilizing *Sox9* transcripts. To examine this hypothesis mRNA decay assays were performed using MBNL1^Fl/Fl^ and MBNL1^Fl/Fl^-Tcf21^iCre^ cardiac fibroblasts. In these assays the decay of *Sox9* and *Mbnl1* mRNA was measured as a function of time exposed to the transcription inhibitor Actinomycin D (**Fig. 6D & E**). Mbnl1 gene expression was measured in this assay to confirm gain (AdMBNL1 treatment) and loss of (Cre treatment) function in the appropriate experimental groups (**Fig. 6E**). Prior to actinomyocin-D treatment MBNL1^Fl/Fl^-Tcf21^iCre^ fibroblasts had 53% less *Sox9* expression relative to MBNL1^Fl/Fl^ controls (time 0, **Fig. 6D**), which accelerated the decay rate of this transcript with transcription inhibition. Adenoviral transfection of MBNL1 (AdMBNL1) stabilized *Sox9* expression in both MBNL1^Fl/Fl^-Tcf21^iCre^ and MBNL1^Fl/Fl^ cardiac fibroblasts, which in turn slowed actinomyocin-D mediated decay (**Fig. 6D**). Together these data suggest that the upregulation of MBNL1 in cardiac fibroblasts following MI acts to stabilize key transcripts like *Sox9* that are critical to state transitions underlying the heart’s fibrotic response.

**Figure 6:**
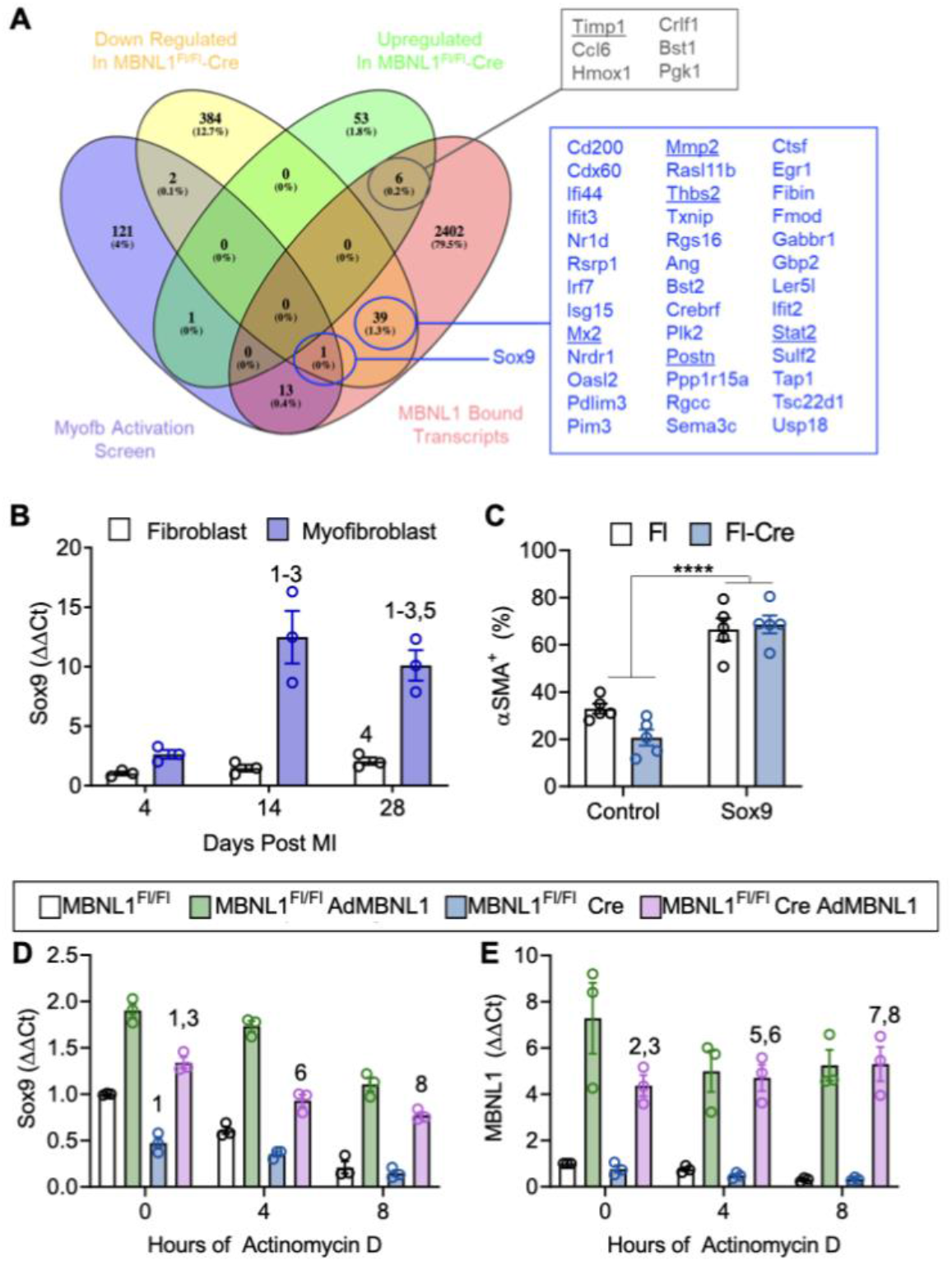
MBNL1 Stabilization of Sox9 promotes cardiac fibroblast to myofibroblast transitions. **(A)** Venn diagram of RNAseq analysis and previously published data showing the overlap between MBNL1 regulated transcripts (pink oval), positive inducers of myofibroblast differentiation (purple oval) (Davis et al., 2015), and genes down regulated in MBNL1^Fl/Fl^-Tcf21^iCre^ cardiac fibroblasts (yellow oval). **(B)** Fold change in *Sox9* gene expression in flow sorted cardiac myofibroblasts (MEFSK4^+^, mG^+^) and fibroblasts (MEFSK4^+^) relative to day 4 fibroblast (MEFSK4^+^) group. Gene expression was normalized to 18s expression and fold change calculated using the ΔΔCt method, dots represent biological replicates, n=3 mice in each condition and bars represent mean±SEM, ANOVA statistical test with Tukey post hoc comparisons. Numbers represent significant pairwise comparisons at p<0.05 **1**-4d Fibroblast, **2**-4d Myofibroblast, **3**-14d Fibroblast, **4**-14d Myofibroblast, **5**-28d Fibroblast, **6**-28d Myofibroblast. **(C)** Quantification of αSMA^+^ stress fiber formation in MBNL1^Fl/Fl^-Tcf21^iCre^ and MBNL1^Fl/Fl^ cardiac fibroblasts transfected with *Sox9.* Dots represent biological replicates, n=5 in each condition and bars represent mean±SEM with ANOVA statistical test with Tukey post hoc comparisons,****p<0.0001 between treatment groups. **(D)** Sox9 and **(E)** MBNL1 gene expression analysis in cardiac fibroblasts from MBNL1^Fl/Fl^-Tcf21^iCre^ and MBNL1^Fl/Fl^ cardiac fibroblasts with or without AdMBNL1 infection and actinomyocin-D treatment. Dots represent biological replicates, n=3 mice in each condition, bars represent mean±SEM and ANOVA statistical test with Tukey post hoc comparisons. Numbers represent significant pairwise comparisons at p<0.05 **1**-0hr Fl, **2**-0hr Fl AdMBNL1, **3**-0hr Fl-Cre, **4**-0hr Fl-Cre AdMBNL1, **5**-4hr AdMBNL1, **6**-4hr Fl-Cre, **7**-8hr Fl Ad BNL1, **8**-8hr Fl-Cre.

## DISCUSSION

The results of this study confirm that MBNL1 plays an essential role in the heart’s fibrotic response through regulating MI-dependent reprogramming of the cardiac fibroblast transcriptome to new cell states needed for the fibrotic phase of cardiac wound healing. This compliments previous work by our lab using global MBNL1 knockout models, which suggested that MBNL1 promotes fibroblast to myofibroblast differentiation through maturing the transcriptome but failed to prove the cardiac fibroblast specific role (Davis et al., 2015). By employing cardiac fibroblast gain and loss of MBNL1 function there is now robust evidence that MBNL1 modulates cardiac fibroblast state transitions to control the degree of fibrotic scarring following MI. By toggling MBNL1 expression we shifted the cardiac fibroblast transcriptome between that of a scar-forming myofibroblast to an earlier progenitor state with gain or loss of MBNL1 function respectively. These data demonstrate that the myofibroblast state and hence fibrotic scarring can be prevented by reverting cardiac fibroblasts to an earlier developmental state (**Fig. 5A**). This provides new evidence that the quiescent adult cardiac fibroblast state is reversible *in vivo* despite the longstanding notion that adult cardiac fibroblast transited to more stably differentiated states following disease or injury. This in turn had additional positive outcomes in terms of reducing maladaptive myocardial remodeling (**Fig. 4C & F**). Notably, blocking a later state transition to that of a matrifibrocyte may also underlie some of these positive outcomes, as a reduction in matrifibrocyte markers were measured in MBNL1^Fl/Fl^-Tcf21^iCre^ cardiac fibroblasts during the late phase of remodeling (**Fig. 4I**). The data presented here, however, cannot discriminate whether MBNL1 directly regulates matrifibrocyte state transitions or whether cardiac fibroblasts must first transition to a myofibroblast before becoming a matrifibrocyte (Fu et al., 2018), and it is this MBNL1-regulated step that actually prevents this additional state transition.

While MBNL1 overexpression has strong effects on a cardiac fibroblast’s transcriptional state at baseline, MBNL1 Tg-Tcf21^iCre^ mice did not have a corresponding induction of cardiac fibrosis as was seen in other mouse models in which overexpression of activators of the myofibroblast state had cardiac fibrosis in the absence of injury (Davis et al., 2015; Molkentin et al., 2017). In this case it is thought that MBNL1 primed the transcriptome by lowering the activation energy needed to transition from a stable quiescent fibroblast state and subsequent exposure to stress like the rigid environment of tissue culture plastic was needed to fully achieve a physiologic myofibroblast phenotype (**Fig. 2L**). To this end it was surprising that a significant increase in scarring and myofibroblast density was not observed in MBNL1 Tg-Tcf21^iCre^ hearts following MI. However, the inability of MBNL1 Tg-Tcf21^iCre^ cardiac fibroblasts to proliferate likely impacted the number of progenitors that could transition to the myofibroblast state (**Fig. E-G**). These data add to a growing body of evidence that molecular profiles alone don’t always emerge at the proteomic or organ physiology level necessitating phenotype validation with physiologic assays (Liu et al., 2016; Ma et al., 2017).

MBNL1 regulates a large number of transcripts that shape a cardiac fibroblast’s molecular state during the injury response with a key mechanism being the stabilization of *Sox9* mRNA. This transcription factor was recently identified as a potential regulator of the fibrotic response through unbiased tomo-sequencing in mouse hearts after ischemia reperfusion injury (Lacraz et al., 2017). *Sox9* has binding sites in key ECM genes needed for the post infarct remodeling response, such as Collagen1a2, Fibronectin1, and Vimentin. *Sox9* expression is relatively low in the mouse heart except during development where it promotes EMT and ECM organization (Akiyama et al., 2004; Lincoln et al., 2007). Recent studies using genetic excision of *Sox9* in myofibroblasts have also demonstrated a role for *Sox9* in the heart’s fibrotic and inflammatory response following MI (Ruiz-Villalba et al., 2020; Scharf et al., 2019). Adding to these findings, it was demonstrated here that the native expression of *Sox9* and *Mbnl1* are greatly enhanced in myofibroblasts following injury and that MBNL1, through its stabilization of *Sox9,* reduces myofibroblast and matrifibrocyte genes similar to findings in *Sox9* knockout mice (Scharf *et al.,* 2019). Collectively, these findings indicate that MBNL1’s role in cardiac fibroblasts is to stabilize core transcripts required for state transitions that underlie the heart’s fibrotic response.

The ability to control myofibroblast reversibility holds great promise for addressing the clinical burden of fibrosis, given that preventing myofibroblast state transitions by pharmacologic or genetic inhibition of its molecular inducers can significantly reduce the fibrotic response in the heart and other tissues (Bugg et al., 2020; Davis et al., 2012, 2015; Dobaczewski et al., 2010; Huang et al., 2019; Koitabashi et al., 2011; Lighthouse et al., 2019; Molkentin et al., 2017; Small et al., 2010). Targeting posttranscriptional mechanisms underlying fibroblast to myofibroblast differentiation is an appealing approach as approximately one-third of the changes in gene expression are subject to translational regulation affecting overall protein abundance (Chothani et al., 2019). MBNL1 has desirable therapeutic qualities, as it regulates multiple facets of fibroblast function that are required for wound healing including contraction and the secretion of ECM that forms a protective fibrotic scar, which in turn could be used to overcome the clinical burden of a permanently scarred heart following stress or MI-injury (Hinz, 2007; Gourdie, Dimmeler and Kohl, 2016).

### Limitations Of The Study

The aim of this study was to understand the fibroblast specific role of MBNL1 in regulating the hearts fibrotic response. Although these data demonstrate that loss of MBNL1 function in cardiac fibroblasts prior to MI prevents fibroblast to myofibroblast state transitions and subsequent fibrotic remodeling, the effects of MBNL1 excision on fibroblast and myofibroblast heterogeneity and fibroblast population dynamics remain unclear. Since cellular heterogeneity is lost when preforming RNAseq, single cell transcriptional analysis would provide additional insights regarding the diversity of cell state changes caused by MBNL1 manipulation following MI. Notably, MBNL1-dependent alterations in fibroblast proliferation could have long lasting effects on the cellular stoichiometry of the heart which in turn could alter cardiac structure and function. Furthermore, direct proof that MBNL1 expression is required for maintaining the myofibroblast state is needed to fully understand the role of MBNL1 myofibroblast state stability and its control over the transition to a matrifibrocyte or quiescent state at the cessation of injury.

## AUTHOR CONTRIBUTIONS

DB, RB, KB, AR, JG, GF, AS, CD and JD conducted experiments and analyzed results. Experiments were designed by DB and JD. DB, CD, and JD contributed to writing and reviewing the manuscript.

## ACKNOWLEDGEMENTS

This work was supported by grants from the National Institutes of Health for JD (HL141187 & HL142624) and CD (R35GM138036). The Biological Mechanisms of Healthy Aging Training Program NIH for DB (T32AG066574). Graduate Research Fellowship from the National Science Foundation for RB (2018261576).

## EXPERIMENTAL PROCEDURES

### Animal Models

All animal experimentation was approved by the University of Washington’s Institutional Animal Care and Use Committee. Lineage reporter mice were generated by crossing a mouse containing a tamoxifen (Tam)-inducible Cre recombinase cassette knocked into the periostin locus (Postn^iCre^) with mice contain a membrane targeted dual fluorescent reporter (mT/mG) knocked into the Rosa26 locus (**Fig. 1F**). All cells in Postn^iCre^-mT/mG mice express membrane-targeted TdTomato (mT) unless Cre is expressed which excises the TdTomato (mT) and moves a membrane-targeted green fluorescent protein (mG) sequence in frame for expression (Kanisicak et al., 2016; Muzumdar et al., 2007). Cardiac fibroblast specific MBNL1 overexpression mice (MBNL1 Tg-Tcf21^iCre^) were generated by crossing mice containing the human MBNL1 cDNA with mice containing a Tam-inducible *Tcf21-Cre* driver (Tcf21^iCre^, (Acharya et al., 2011, 2012; Davis et al., 2015)). Conditional MBNL1 knockout mice (MBNL1^Fl/Fl^) were generated using targeted C57BL/6 embryonic stem cells (ES) from the International Knockout Mouse Consortium (IKMC). Founders were bred onto a C57BL/6 background and then to mice expressing Flippase to excise the LacZ-neomycin cassettes still present within the floxed MBNL1 allele, as IKMC uses a knockout first approach for their targeted alleles. Once the LacZ-neomycin cassettes were flipped-out, MBNL1^Fl/Fl^ mice were crossed with Tcf21^iCre^ to generate cardiac fibroblast specific MBNL1 knockout mice. Tamoxifen induction of Cre recombinase expression was started between 6-8 weeks of age and achieved with 5 days of intraperitoneal (IP) injections of pharmaceutical grade tamoxifen dissolved in peanut oil (25mg/kg) followed by 9 additional days on tamoxifen citrate chow (400mg/kg body weight, Harlan Laboratories). Mice remained on Tam chow until the experimental end point unless stated otherwise.

### Surgical Model of Myocardial Infarction (MI)

The surgical model was previously described (Bugg et al., 2020; Molkentin et al., 2017), but briefly 8-week-old mice were anesthetized using injectable ketamine and xylazine. Mice were mechanically ventilated through oral intubation and a lateral thoracotomy was performed to expose the left ventricle. The pericardium was removed, and the left anterior descending artery was permanently ligated using 8-0 Surgipro tapered suture. Two days prior to surgery mice were taken off Tam chow for 2 days to prevent any adverse effects of Tam during the surgical procedure and then put back on Tam chow until the studies end point. At harvest hearts were excised, rinsed in phosphate buffered saline (PBS), and relaxed in saturated potassium chloride solution before being fixed in formalin and prepped for paraffin sectioning. Experimentalists remained blinded to the genotypes until analysis was complete. Both male and female mice were used in all experiments and mice were randomly assigned to groups. Echocardiography was performed on a Vevo2100 under isoflurane anesthetic.

### *In vivo* Proliferation Following MI

Mice underwent MI procedure as stated above. Two boluses of EdU (100mg/kg) were injected IP at 24 hours and 9 hours before harvesting. Mice were then euthanized, and hearts fixed in 4% paraformaldehyde overnight. Tissues were then processed through a sucrose gradient, embedded in optimal cutting temperature compound (OCT), and prepared for 5μm cryosectioning.

### Primary Cardiac Fibroblast Isolation

Cardiac fibroblasts were freshly isolated by Langendorff perfusion with type II collagenase (2mg/ml) and liberase blendzyme (0.4mg/ml) solubilized in Krebs-Henseleit buffer as previously described (Bugg et al., 2020; Molkentin et al., 2017). For culture experiments cardiac fibroblasts were plated in in Dulbecco’s Minimal Essential Media (DMEM) with high glucose and supplemented with 1% penicillin and streptomycin (P/S), and 20% fetal bovine serum (FBS). Cardiac fibroblasts were expanded to passages 3-5 for experimentation. For some experiments cell-permeant Cre recombinase (TAT-Cre,1:100, EMD Millipore) was added to the cultures for 2 days to permanently excise MBNL1. For differentiation assays cells were cultured in differentiation media (DMEM + 1% FBS + 1% P/S) with or without recombinant TGFβ (R&D systems, 10 ng/mL) and analyzed 72 hours later by immunofluorescent staining described below. For proliferation assays cardiac fibroblasts were treated with 10μM EdU every 12 hours over 1 day in 2% FBS media. At the studies end point cardiac fibroblasts were fixed in 4% PFA and Click-iT chemistry (Invitrogen) used to detect EdU positivity per the manufacturer’s instructions.

### TGFβ Treatments, Adenoviral Gene Transfer & Sox9 Overexpression

Recombinant TGFβ (10 ng/ml, R&D System) was used to induce myofibroblast transformation. For experiments that used adenoviral gene transfer, cardiac fibroblasts treated with adenovirus overnight followed by a media change and cells examined 3 days after induction. The following adenoviruses have been previously described: ΔCnA, SRF, GFP, and MBNL1 (Davis et al., 2012, 2015; Liu et al., 2001; Wilkins et al., 2004). An adenovirus expressing GFP was used as a control. Sox9 overexpression was obtained by transfecting cardiac fibroblasts with full length Sox9 cDNA from the mammalian genome collection (Dharmacon) using X-treme Gene transfection reagent (Sigma) diluted in Opti-MEM media at a 4:1 ratio of transfection reagent to plasmid. βgal plasmid was used for control transfections.

### Collagen Gel Contraction Assay

Cardiac fibroblasts were isolated, expanded and treated with adenovirus or TGFβ 24 hours prior to seeding into collagen gels as previously described (Davis et al., 2012; Ngo et al., 2006). Here 50,000 cardiac fibroblasts were seeded into each gel and once solidified were released from the plate into differentiation media. Gels were photographed and measured every 24 hours over 4 days. ImageJ software (NIH) was used to calculate the surface area, which are reported as values normalized to the initial size of the gel. Data shown is at 48 hours post seeding.

### Transcript Stability Assay

Tam-treated MBNL1^Fl/Fl^ and MBNL1^Fl/Fl^-Tcf21^iCre^ cardiac fibroblasts were infected with either AdGFP (controls) or AdMBNL1 for 36 hours at which time transcription was inhibited with 2μg/ml Actinomycin-D (Sigma). RNA was isolated from each group after 0, 4, and 8 hours of treatment, reverse transcribed, and analyzed by real-time PCR as previously described (Davis et al., 2015).

### Cardiac Fibroblast & Myofibroblast Purification by Flow Cytometry

For Postn^iCre^-mT/mG gene expression and RNA sequencing experiments cardiac fibroblasts were isolated and the subjected to flow cytometry as previously described (Molkentin et al., 2017). Briefly cells were strained through a 70 μm cell strainer and then stained with CD11b (1:50 Miltenyi Biotec 130-113-800) and MEFSK4 (1:50 Miltenyi Biotec 130-120-802). This allows for the removal of CD11b cells which are known to express high levels of MBNL1. Cells were then sorted on an Aria II live cell sorter. For Postn^iCre^-mT/mG experiments the same antibody scheme was used but the MEFSK4^+^ GFP^-^ and MEFSK4^+^ GFP^+^ populations were divided so that analysis could be done on both populations from the same heart. RNA isolation was then preformed using an RNAqueous Micro Kit which is specifically designed for low RNA yields. Gene specific amplification was used during cDNA synthesis to increase signal of target genes.

### Gene expression Analysis

Real time polymerase chain reaction (RT-PCR) methods are previously described from our laboratory (Bugg et al., 2020; Davis et al., 2012, 2015; Molkentin et al., 2017). Briefly, total RNA was extracted using RNAqueous Micro Kit for all flow sorted samples or using QIA shredder homogenization and the Qiagen RNeasy kit for in vitro culture. Total RNA was reverse transcribed into cDNA using random hexamer primers and SuperScript III first-strand synthesis kit (Invitrogen) according to the manufacturer’s instructions. RT-PCR was performed on a CFX96 Real-Time System with a Biorad C1000 Touch Thermal Cycler using Sso Advanced SYBR Green (Biorad). Thermocycler conditions were as follows: Polymerase Activation and DNA Denaturation at 95°C for 30s, Denaturation at 95°C for 5s, Annealing/Extension and Plate Read at 56°C for 30s. 39 cycles of denaturation and annealing were performed. Fold changed in gene expression was determined using the 2^ΔΔCT^ method. Any differences in cDNA were correct by calculating the difference (ΔCT) between the target gene’s threshold cycle (CT) and the CT for 18s, which serves as the housekeeping gene. Primer sequences are as follows: Acta2, Fwd: ACCCACCCAGAGTGGAGAAG, Rev: AGCATCATCACCAGCGAAG; Col1a1, Fwd: AATGCAATGAAGAACTGGACTG, Rev: CCCTCGACTCCTACATCTTCTG; MBNL1 (Mouse), Fwd: CACTGAAAGGTCGTTGCTCCA, Rev: CGCCCATTTATCTCTAACTGTGT; MBNL1 (Human) Fwd:CTCTGTCCGGTTGACAGGC, Rev:CTGAAAACATTGGCACGGGT; Sox9, Fwd:TATCTTCAAGGCGCTGCAA, Rev: TCGGTTTTGGGAGTGGTG.

### RNAseq Analysis

Cleaned RNAseq reads were uploaded to the public Galaxy server at usegalaxy.org for bioinformatic analysis (Afgan et al., 2018). Reads were aligned using HISAT2 to the mm10 reference genome, summarized using featurecounts, and differential expression between timepoints and genotypes were tested using DESeq2 (Kim et al., 2019; Liao et al., 2014; Love et al., 2014). Genes with an adjusted p < 0.05 were considered significantly differentially expressed. PCA was performed on the log2 normalized counts matrix from DESeq2 using the scikit-learn package (Pedregosa et al., 2011). Heatmaps and gene expression scatterplots were generated from the log2 normalized counts output from DESeq2 using the seaborn package in Python (Waskom et al., 2020). Functional enrichment analysis was conducted using the GProfiler web app, inputting all significant differentially expressed genes as an ordered query (Raudvere et al., 2019).

To assess time course expression of clustered, a pairwise correlation matrix between time course-significant genes was calculated, hierarchically clustered using the scipy package in Python, and plotted with seaborn.(Virtanen et al., 2020; Waskom et al., 2020) Functional enrichment analysis on genes from each cluster was performed using GProfiler, and log2 normalized gene expression levels per cluster were calculated and plotted over time using seaborn (Raudvere et al., 2019).

### Statistical analysis

Time course plots (**Fig. 1C-E**) were generated with 95% confidence intervals using the pointplot function in seaborn (Waskom et al., 2020). Prism versions 7 and 8 were used for plotting all other data and statistical analysis. Data are represented as mean ± SEM. 2-way ANOVA with Tukey post hoc analysis was used for multiple comparisons. Two-tailed t-tests were used for pairwise comparisons with a p < 0.05 considered significant.

### Western Blot

Tam-induced MBNL1^Fl/Fl^ and MBNL1^Fl/Fl^-Tcf21^iCre^ fibroblasts treated with 10ng/ml TGFβ and cell lysates were collected 48 hours later in RIPA buffer [5 M NaCl, 10% Triton-X 100, 25%SDS, 1 M Tris-Cl PH 7.4]. Lysates were diluted in Laemmli buffer, 20μg of protein was loaded into 10% SDS-PAGE acrylamide gels and transferred to PVDF membrane for immunodetection. MBNL1 was detected with anti-MBNL1 (1:100, rabbit polyclonal antibody, Abcam) overnight at 4°C with goat anti-rabbit HRP conjugate secondary at 1:10,000 from Santacruz biotechnology for 1 hour. Anti-GAPDH (1:10,000, mouse monoclonal, Fitzgerald Industries) overnight at 4°C with goat anti-mouse HRP conjugate at 1:10,000 from Santacruz for 1 hour was used as a loading control.

### Staining procedure *in vivo*

Hearts were cut in half on the transverse plane prior to processing and sectioning. 5μm paraffin sections were obtained for Sirius Red/Fast Green staining. This method stains muscle tissue in green and fibrotic scar in red. Images of whole hearts were taken at 2x magnification and quantified in imageJ using color thresholding. Serial sections were then used for αSMA (1:500 Sigma) and IsolectinB4 (IB4) (10ug/mL Vector Biolabs) staining t o quantify myofibroblast number. These methods are previously described from our laboratory, but briefly sections were deparaffinized and the blocked in PBS with 1 % BSA and 0.1 % cold fish skin gelatin. Primary antibodies were incubated overnight in blocking solution at 4°C(Bugg et al., 2020; Molkentin et al., 2017). AlexaFluor secondary antibodies (1:1000 Invitrogen) were used for 1.5 hours at room temperature to detect the antigen. Hoechst (1:2000 Thermo Fisher) was used to visualize Nuclei. Additional groups were incubated in primary or secondary antibody alone to control for non-specific signaling for imaging analysis. To visualize EdU staining in vivo Click-iT chemistry was used following the manufactures instructions with the substitution of normal horse serum (NHS) for PDGFRα co-staining.

### Staining procedure *in vitro*

Immunofluorescence staining procedures followed previously described methods from our laboratory (Bugg et al., 2020; Molkentin et al., 2017). Briefly, cardiac fibroblasts were fixed in 4% paraformaldehyde, permeabilization in PBS containing 0.1% Triton-X100, and blocked in PBS containing 0.1% Triton-X100 and 10% Normal Goat Serum (NGS). Primary antibody for αSMA (1:500 Sigma) was incubated overnight at 4 °C. AlexaFluor secondary antibodies (1:1000 Invitrogen) were used for 1.5 hours at room temperature to detect the antigen. Hoechst was added with secondary antibody used to visualize nuclei. Additional groups were incubated in primary or secondary antibody alone to control for non-specific signaling for imaging analysis.

### Image analysis

To quantify αSMA^+^ and EdU^+^ fibroblasts *in vitro,* cells were manually counted across 10 representative fields of view (FOVs) and normalized to the total number of nuclei to get a percent of total population. To score the number of αSMA^+^ myofibroblasts in baseline MBNL1 Tg-Tcf21^iCre^ mice and NTG-Tcf21^iCre^ controls, the entire cross section of a heart was imaged and the total number of αSMA^+^IB4^-^ cells were manually scored. Since there are so few cells in these hearts they are shown as raw values and not normalized to fibroblast number or heart area. For infarcted tissue 9-16 FOV of the infarct and border zone were imaged and the number of αSMA^+^IB4^-^ cells were scored. This was then normalized to the average number of cells per FOV. For in vivo proliferation quantification we first scored for the total number of PDGFRα^+^ fibroblasts in the infarct and borderzone using the Cell Counter Plugin in Fiji (NIH). Then the Click-iT staining was overlaid with the PDGFRα signal and we scored for the number of PDGFRα^+^EdU^+^ allowing us to calculate the percent of the total fibroblast population that was proliferating.

