## Supplemental Figure 1 for "Fibroblast State Reversal By MBNL1-Dependent Transcriptome Modification Regulates Cardiac Repair"

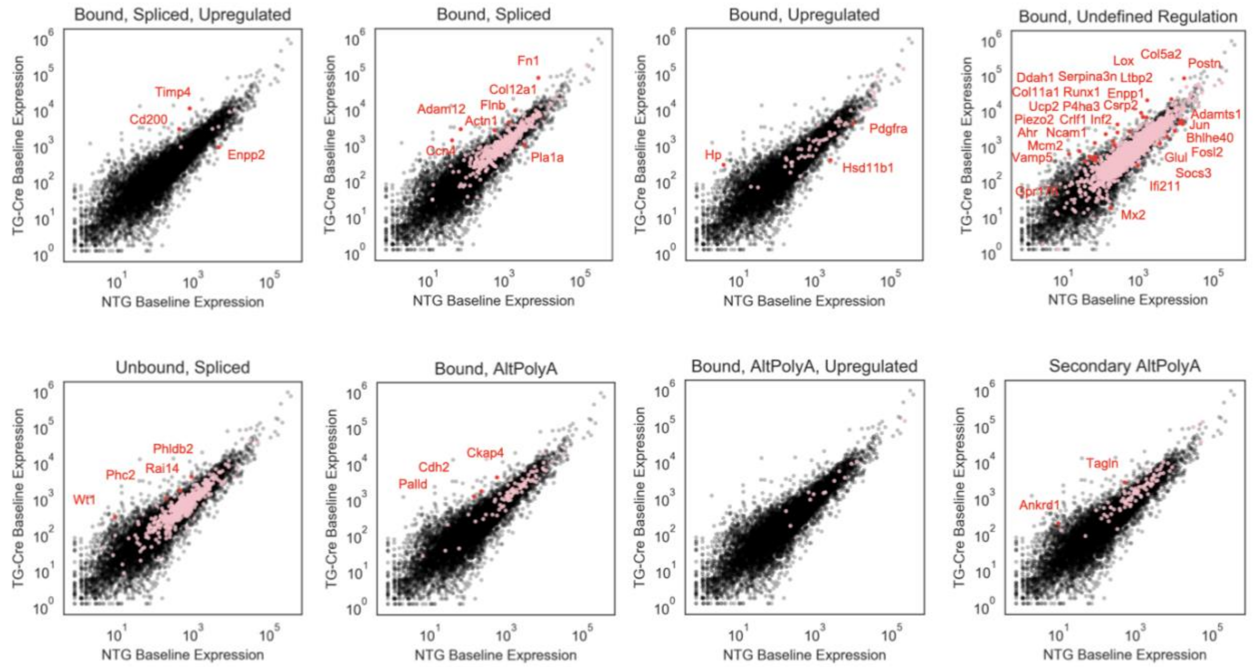


**Supplemental Figure 1:**

Scatterplot of differentially expressed genes between MEFSK4 purified cardiac fibroblasts from MBNL1 Tg-Tcf21$^{iCre}$ (TG-Cre) versus NTG-Tcf21$^{iCre}$ (NTG) in which transcripts that are bound and regulated by MBNL1 are labeled in red and relationships categorized by the type of MBNL1-mediated transcriptional regulation (spliced, stability, or differential polyadenylation). Axes represent the normalized gene expression levels in wild type and MBNL1 overexpressing fibroblasts.
